# Oleoylethanolamide effects on stress-induced alcohol consumption: a lipid at crossroads between stress, reward and neuroinflammation

**DOI:** 10.1101/2023.11.06.565786

**Authors:** Macarena González-Portilla, Sandra Montagud-Romero, Susana Mellado, Fernando Rodríguez de Fonseca, María Pascual, Marta Rodríguez-Arias

**Author notes:** Correspondence author: Dr. Marta Rodríguez-Arias. Department of Psychobiology, Facultad de Psicología, Universitat de Valencia, Avda. Blasco Ibáñez, 21, 46010 Valencia, Spain.

## Abstract

The endocannabinoid system is involved in multiple drug-related behavior as well as in the stress response. The transient increase in endogenous cannabinoids as well as endocannabinoid-like molecules contributes to healthy adaptation to stress exposure. In this study, we tested the effect of systemic OEA treatment (10mg/kg) before or after social defeat (SD) on alcohol self-administration (SA). Mice were divided into non-stressed (EXP) and stressed mice (SD) and randomly assigned to a treatment condition (CTRL, OEA or 10OEA). Mice in the EXP/SD-OEA group received four doses before each SD encounter while mice in the EXP/SD-10OEA mice received 10 daily doses after stress exposure. Three weeks after SD, mice were trained to alcohol 20% (vol/vol) SA. Upon extinction, a cue-induced reinstatement test was performed. Our results showed that only multiple-dose chronic OEA treatment (SD-10OEA group) was effective in preventing the stress-induced increase in alcohol consumption observed in defeated mice. We did not observe any effects of OEA on relapse-like behavior. Altogether, these data suggest that exogenously increasing OEA levels counteracts the adverse effects of stress on alcohol drinking.

## 1. Introduction

Stress plays a fundamental role for the development and trajectory of alcohol use disorder (AUD). AUD patients present an increased risk of developing comorbid mental disorders (Castillo-Carniglia et al., 2019). Individuals may seek for alcohol to alleviate depressive and anxiety symptomatology. In turn, as a result of alcohol abuse, AUD patients experience negative mood affect stemming from withdrawal, feelings of craving, frustrated abstinence and the alcohol-related negative consequences on social functioning. A great effort is being made to identify pharmacological agents that aim to restore the altered stress system, target reward dysfunction and improve negative mood symptomatology in AUD (Burnette et al., 2022).

In the past years, signaling lipids have been shown to be important modulators of the stress response. N-acylethanolamines are a family of fatty acid amides (anandamide, AEA; oleoylethanolamide, OEA; palmitoylethanolamide, PEA) that interacts with the endocannabinoid (eCS) system), (Tsuboi et al., 2018). Unsaturated AEA has activity on cannabinoid (CB)1 receptors while unsaturated OEA and PEA are considered endocannabinoid-like congeners that indirectly modulate eCS function. In the periphery and in the brain, NAEs are present at low concentrations but are also quickly synthesized from lipid substrates in cellular membranes in response to various types of physiological demands, including stress.

Converging evidence from animal and human research demonstrates that the increase in NAEs after acute stress exposure modulates the hypothalamic-pituitary-adrenal (HPA) axis, contributes to adaptation and limits the adverse consequences of the stress response (Gorzalka et al., 2008; Hill et al., 2010). For instance, AEA serum concentrations have been revealed to function as a protector factor for PTSD in humans exposed to trauma (Mayo et al., 2022). Also, increased peripheral biomarkers of eCSs function have been associated to resilience to substance use disorder after childhood maltreatment (Perini et al., 2023). In the laboratory, healthy participants exposed to a stress task exhibited elevated levels of AEA and OEA (Dlugos et al., 2012). In this sense, an active eCBs tone has been associated to a dampening of the physiological adverse effects of stress (deRoon-Cassini et al., 2020).

In preclinical research, pharmacological boosting the eCS signaling can improve depressive-like and anxious behavior (Parolaro et al., 2010). Systemic administration of OEA has been proven to have antidepressant activity using models of physical (Costa et al., 2018; Jin et al., 2015) and social stress (Rani et al., 2021). Furthermore, OEA treatment before SD prevented the SD-induces potentiation of the rewarding properties of cocaine (González-Portilla et al., 2021).

Interestingly, OEA has also inhibitory effects on alcohol consumption. OEA treatment (10mg/kg) reduces alcohol self-administration (SA) and consumption (Sánchez-Marín et al., 2022, Bilbao et al., 2016; Gonzalez-Portilla, *in preparation*).

Targeting the adverse effects of stress is a useful strategy for the treatment of AUD since many symptoms can be considered to originate from a maladaptive stress response. Therefore, considering the stress-buffering of OEA and its modulating effects on alcohol consumption, we hypothesize that OEA could be particularly effective in mitigating the stress-induced alcohol SA. In this study, we tested the effects of different OEA treatments in modulating alcohol SA in socially defeated mice.

## 2. Methods and materials

### 2.1 Experimental design

OF1-strain male mice (Charles River, France) used in this study were housed in groups of four in standard plastic cages. Mice were under constant temperature and a reverse 12-h light/dark cycle (lights on at 7:00h) and provided with food and water *ad libitum*, except during behavioral testing. All procedures were conducted in compliance with the guidelines of the European Council Directive 2010/63/EU regulating animal research and were approved by the local ethics committees of the University of Valencia (__).

### 2.2 Drug administration

OEA was synthesized as described in (Rodríguez De Fonseca et al., 2001). OEA was dissolved in 5% Tween 80 in saline and injected by intra peritoneum route (IP, 10mg/kg) 10 minutes before the corresponding timepoint according to the experimental condition. The doses were chosen according to previous studies in mice reporting effective therapeutic and anti-inflammatory effects (González-Portilla et al., 2023; Moya et al., 2021; Rivera et al., 2018; Sayd et al., 2015).

**Figure 1.**
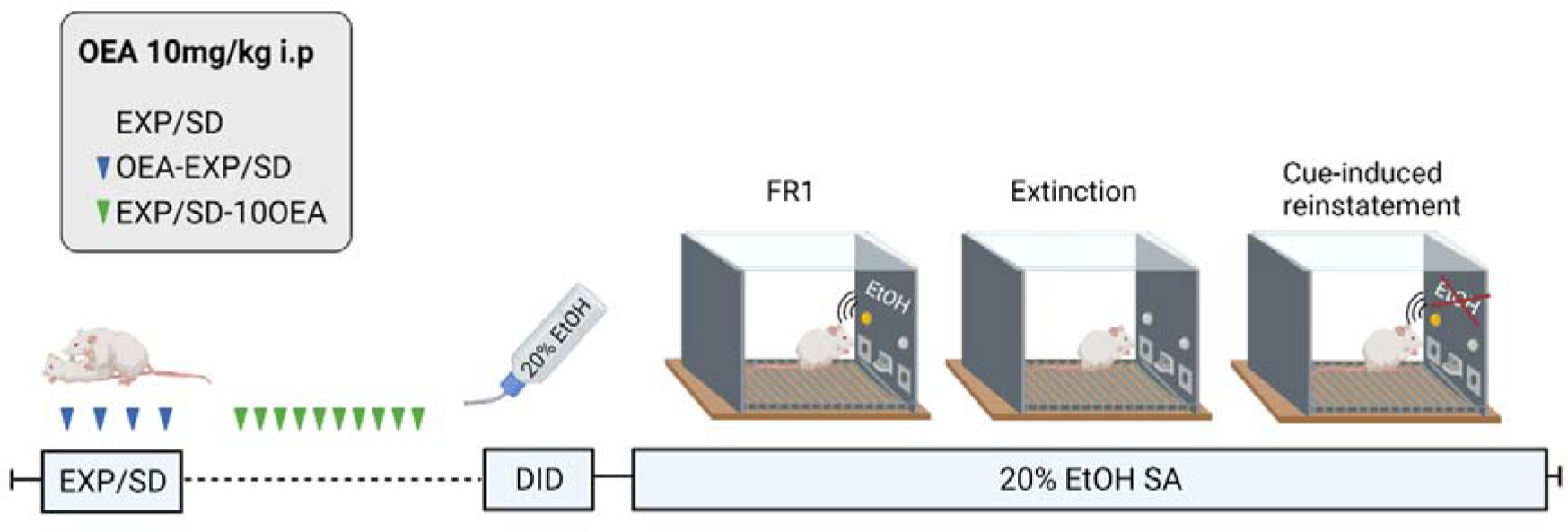
Experimental design. Mice were randomly assigned to a stress condition (exploration, EXP; social defeat, SD) and treatment (EXP/SD-OEA or EXP/SD-10OEA). After three weeks from the stress procedure, mice were habituated to alcohol with the Drinking in the Dark test. Next, mice were trained to self-administer alcohol 20% (vol/vol) on a fixed-ratio 1 (FR1). Upon extinction, mice were tested on a single cue-induced reinstatement session.

### 2.3 Social defeat stress

Mice in the stress condition (SD) were exposed to four episodes of SD. In this study, the SD procedure was performed as described in previous works (Rodríguez-Arias et al., 2018). Each of the SD encounters lasted 25 minutes and consisted of three phases, each of which began by introducing the experimental animal into the home cage of the “resident” (a male aggressive opponent). During this first phase, the experimental mouse is protected from the resident by a wire mesh that allows for social interaction and facilitates instigation and provocation. After 10 minutes, the wire mesh is removed to allow confrontation between the two animals for a video-recorded 5-minute period. In the third phase, the wire mesh is set again for further 10 minutes. The non-stressed exploration (EXP) group underwent the same protocol but without the presence of a “resident” opponent in the cage.

### 2.4 Drinking in the Dark

The Drinking in the Dark (DiD) paradigm was employed as a pre-exposure procedure to habituate mice to ethanol before starting the oral SA. Based on the basic paradigm of Rhodes et al. (2005), this voluntary drinking protocol consisted of two phases. On the first day mice were moved from their homecage for two hours to habituate to the individual cages and the drinking tubes. In the second phase of the protocol, mice received 3 days of 2h/day access to 20% (v/v) ethanol solution starting 3h after lights off. On day 4, the procedure extended for 4 h. After the each DiD session, mice returned to their homecage and liquid consumption was recorded immediately after.

### 2.5 Oral alcohol self-administration (SA)

Mice were trained to orally self-administer alcohol (20% vol/vol) during operant sessions of 60-minute duration using procedures previously described (CITAS). The procedure consisted of five phases: training, fixed ratio 1 (FR1), progressive ratio (PR), extinction, and reinstatement. No food or water deprivation was performed.

Voluntary oral ethanol SA administration was assessed in eight modular operant chambers (MED Associated Inc., Georgia, VT, USA) equipped with a chamber light, two nose-poke holes, one fluid receptacle, one syringe pump, one stimulus light and one buzzer. The chambers were placed inside sound-attenuated cubicles. Designated active nose-pokes delivered 36μl of fluid associated to a 0.5s light cue and a 0.5s buzzer beep, which was followed by a 6s time-out period. Inactive nose-pokes triggered no event. Software package (Cibertec, SA, Spain) controlled stimulus and fluid delivery and recorded operant responses.

#### Training (15 days)

Mice were trained to poke to the active hole for 20% (v/v) ethanol solution delivery (20 μl) under a fixed ratio 1 (FR1). Inclusion criteria for the next step in the procedure comprised 60 % discrimination for active, over inactive nose-poke responding across the three last days of this phase.

#### FR1 (10 days)

The number of effective responses and 20% ethanol (v/v) consumption (μl) were measured under a FR1 for 10 daily sessions. After each session, the alcohol solution that remained in the receptacle was collected and measured with a micropipette.

#### PR (1 day)

Mice undertook a single 2h long PR session in which the response requirement necessary to obtain one reinforcement escalated according to the following series: 1-2-3-5-12-18-27-40-60-90-135-200-300-450-675-1000. Breaking point (BP), defined as the highest number of nose-pokes each mouse performed to earn one reinforcement, was used to quantify motivational strength.

#### Extinction sessions

All mice progressed to the extinction phase, which consisted of removal of the cue light, buzzer, and the reward (alcohol) delivery after a response on the active nose-poke during the 60-minute session.

The extinction phase continued until the average in each experimental condition reached the criterion (at least 60% decrease in active lever response for at least three consecutive days).

#### Cue-induced reinstatement (1 day)

Following the extinction phase, a 60-minute single session of cue-induced reinstatement was performed. Effective responses on the active nose-poke were followed by presentation of cue light and buzzer stimuli in the absence of alcohol delivery. For this session, fluid receptacle was primed with alcohol solution.

### 2.6 Tissue sampling and biochemical analyses

Mice were sacrificed by cervical dislocation. The__and__were precisely dissected out based on the atlas of the Paxinos and Franklin using a coronal brain matrix. Tissue samples were stored at -80ºC until the

### 2.7 Statistical analyses

Data relating to body weight and alcohol SA were analyzed by a mixed ANOVA with two between-subjects variable “Stress” with two levels (EXP, SD) and “Treatment” with 3 levels (CTRL, OEA, 10OEA).

A one-way ANOVA with two between-subject’s variable “Stress” with two levels (EXP, SD) and “Treatment” with 3 levels (CTRL, OEA, 10OEA) was employed to analyze alcohol consumption and breaking point values during the PR session.

To test for the cue-induced reinstatement, the number of active nose poke responses were analyzed with a two-way ANOVA with two levels (EXP, SD) and “Treatment” with 3 levels (CTRL, OEA, 10OEA)– and one within subjects’ variable with two levels (Day: extinction and cue-induced reinstatement). Data were analyzed using SPSS Statistics v23. All data are presented as the mean (±SEM). Bonferroni post-hoc tests were also analyzed. Statistical analyses were performed using SPSS Statistics v23.

## 3. Results

### 3.1 Body weight

Mice body weight was measured weekly. The ANOVA revealed a significant effect of the variable week [F(12,792)=5.176, (p<0.001)], as mice exhibited increased bodyweight across weeks compared to the first measurement (p<0.001). No differences were found between groups.

**Figure 2.**
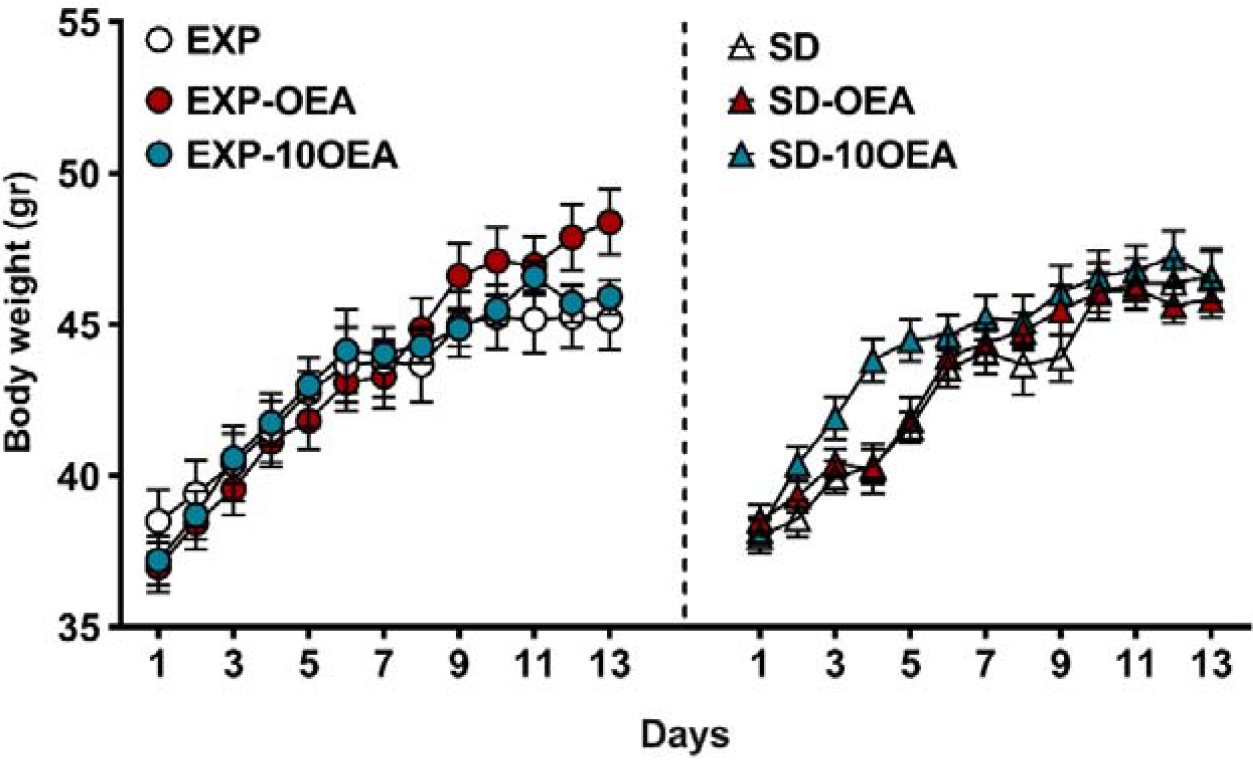
Body weight of mice over 13 weeks. Data are presented as mean (±SEM) amount of body weight (gr).

### 3.2 Alcohol self-administration

The ANOVA for the alcohol consumptionduring the FR1 schedule revealed a significant effect of the variable Day [F(9,531) = 4.970; p < 0.001], with a higher alcohol consumption in Day 2, 3 and 4 compared to Day 7 (p <0.05) and Day 9 (p <0.05). The ANOVA also showed an effect of the interaction Stress x Treatment [F(2,59) = 3.246; p = 0.046]. Non-treated SD mice (SD) and those treated with OEA before each SD (SD-OEA) exhibited increased alcohol consumption compared to their corresponding non-stressed mice (p < 0.001 and p = 0.05, respectively).

In addition, mice SD-10OEA exhibit decreased alcohol consumption compared to non-treated stressed mice (SD) and mice treated with OEA before SD (OEA-SD group), (p<0.001 and p > 0.05, respectively).

Regarding active nose poke responses, the ANOVA revealed a significant effect of the variable MStress [F(1,58) = 6.189; p = 0.016]. SD mice exhibited more active nose poke responses compared to EXP mice (p = 0.016).

**Figure 2.**
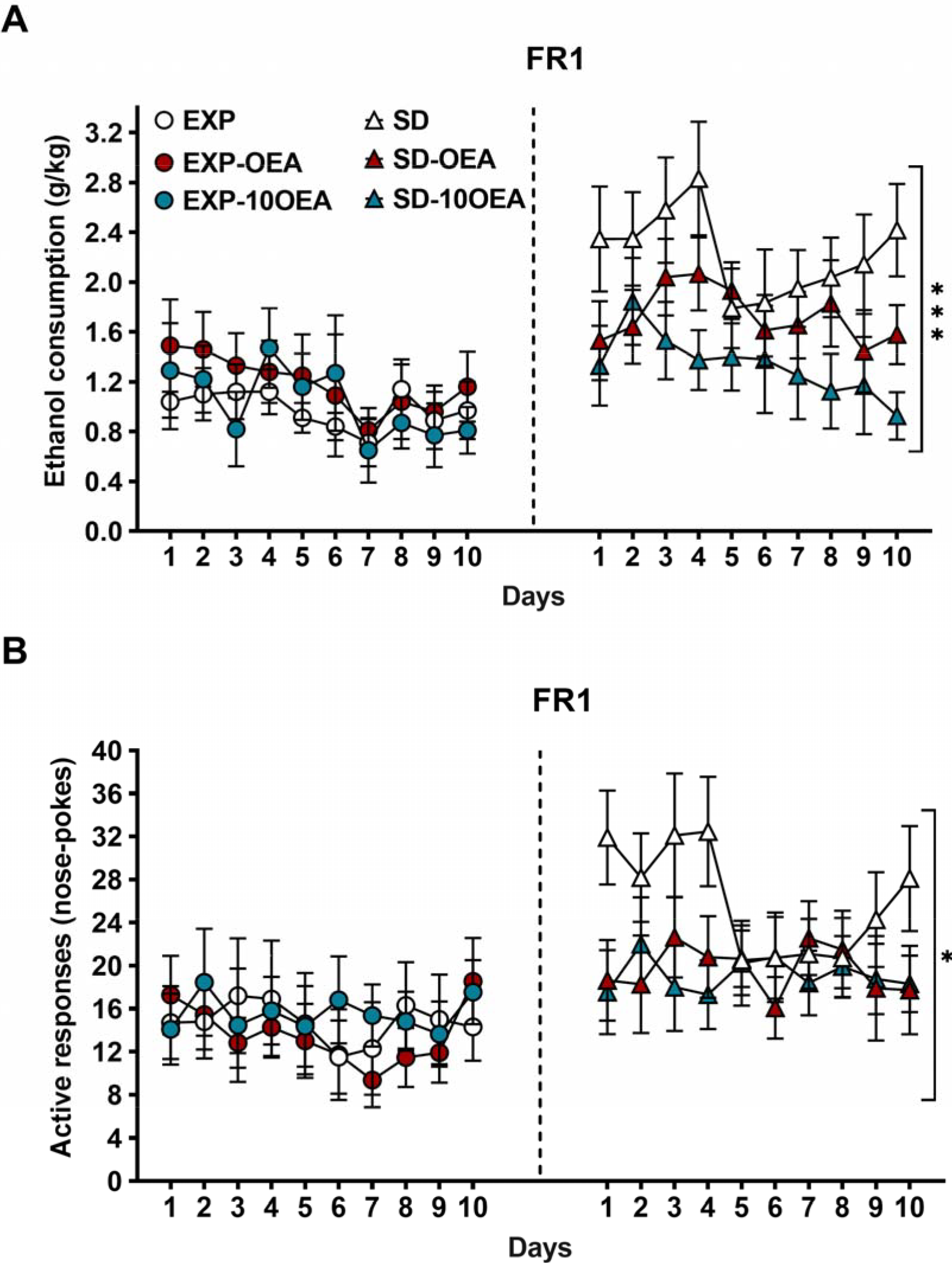
A) Alcohol consumption (g/kg) and B) active nose pokes responses across ten days of operant alcohol self-administration 20% (vol/vol) on a fixed-ratio 1 (mean ± SEM). ***p□<□0.001, *p□<□0.05, significant difference of the SD groups compared to the EXP group.

During the PR, the ANOVA revealed a significant effect of the variable “Stress” [F(5,63)=6.863 (p = 0.011). Post-hoc comparisons showed that breaking point values were increased in SD mice compared to EXP mice. Regarding alcohol consumption, no differences were found between groups.

**Figure 2.**
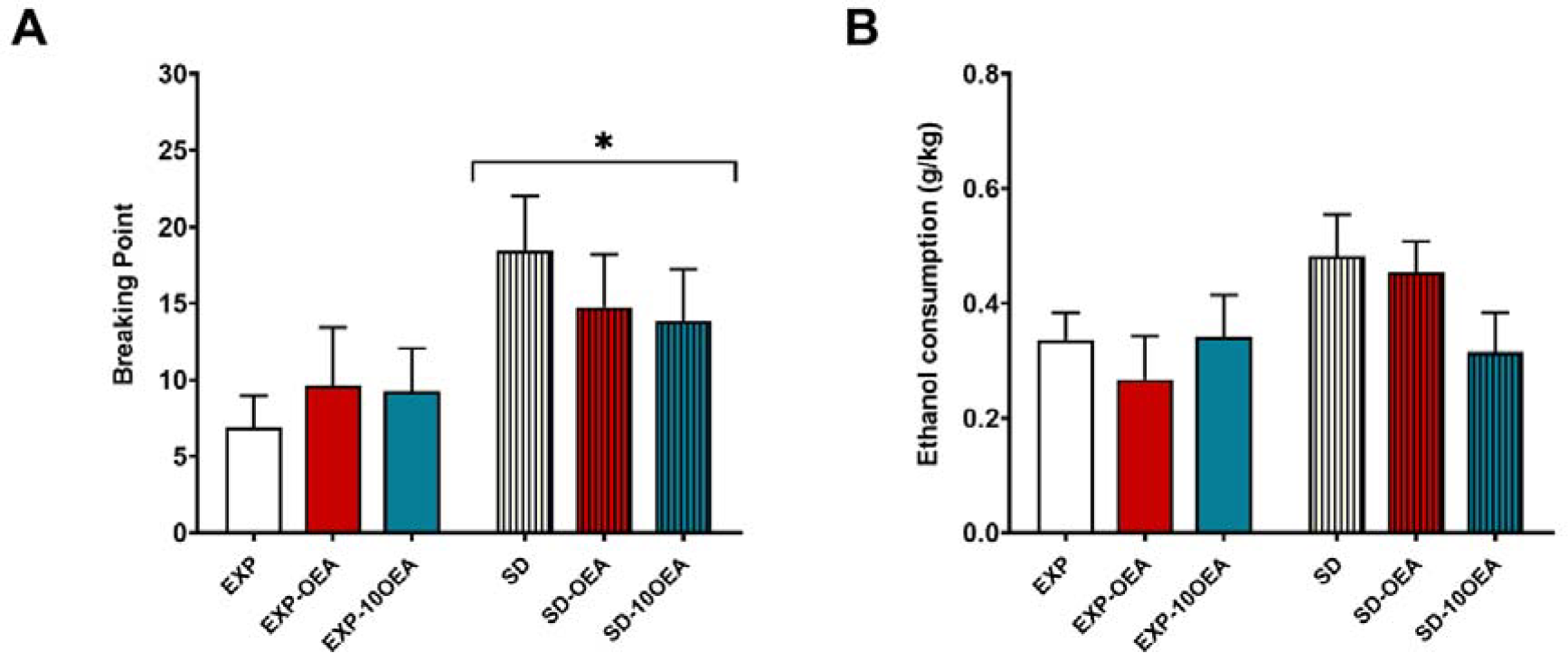
Progressive ratio. The columns represent means and the vertical lines ± SEM of the breaking point values and (d) EtOH consumption during the PR session. *p□<□0.05, significant difference of the SD groups compared to the EXP group.

The ANOVA for the reinstatement test revealed a significant effect of the variable Days [F(1,67)□=□19.034, p□<□0.001]. Active nose poke responses during the reinstatement day were higher than during the extinction (p < 0.001). Increased active nose poke responding during the cue-induced reinstatement test did not significantly differ as a function of OEA treatment.

**Figure 3.**
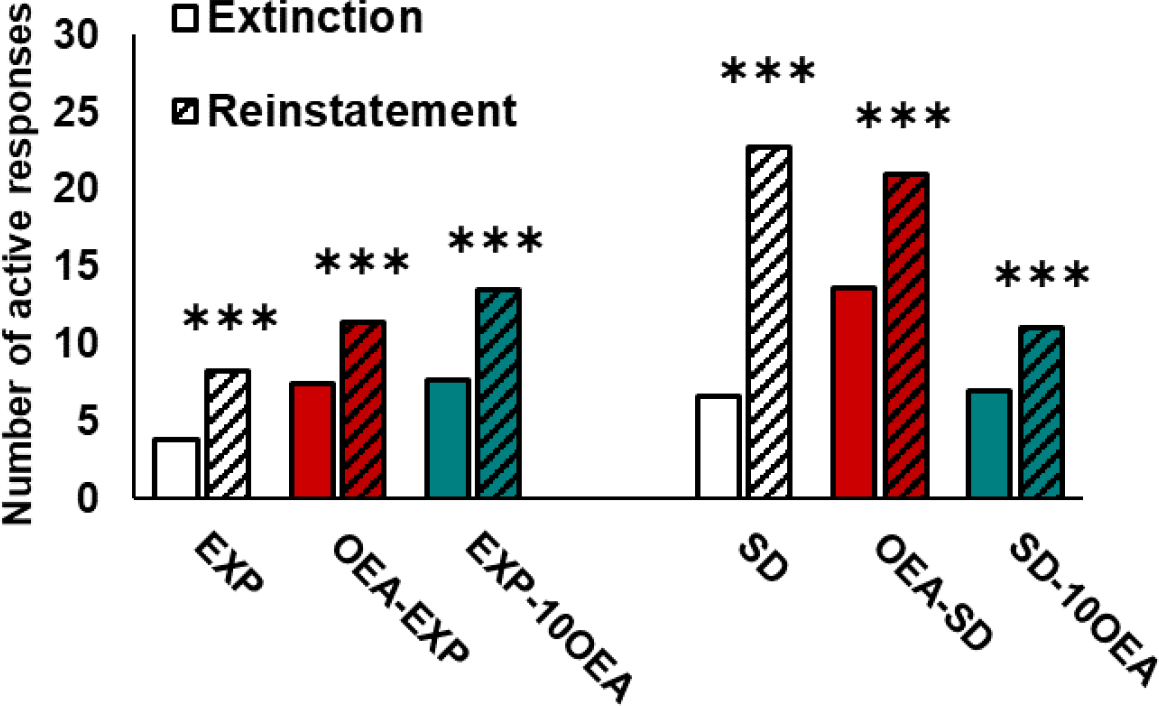
Cue-induced reinstatement. Active and inactive nosepokes responses during extinction and the cue-induced reinstatement test. ***p□<□0.001 Significantly difference from average last 3 days of extinction.

## 4. Discussion

The present study explored how exogenously boosting OEA levels before or after stress exposure influences alcohol consumption in the SA paradigm. Our results confirmed that intermittent SD results in increased in alcohol intake. To test the effects of systemic OEA treatment on alcohol SA we employed two different administration schedules. We observed that a multiple-dose chronic OEA treatment was effective in reducing alcohol consumption during the FR1 period. Additionally, we also studied the long-lasting effects of stress and OEA treatment in a cue-induced reinstatement test.

Regarding drugs of abuse, traumatic experiences and social stress are among the most prominent risk factors for developing AUD (Kendler et al., 1999). SD is a potent stressor with a long-term impact on behavior (Montagud-Romero et al., 2018). Mice exposed to the experience of SD exhibit a well-characterized phenotype involving social withdrawal, anxious and depressive-like behavior (Golden et al., 2011). Concerning drugs of abuse, SD mice exhibit increased vulnerability to the rewarding properties of alcohol and cocaine (Covington & Miczek, 2001; Montagud-Romero et al., 2020). In the SA, SD mice drink more alcohol and work harder to obtain alcohol (Reguilón et al., 2022). In our study, we confirmed the long-lasting effects of social stress as socially defeated mice consumed more alcohol and proved an increased motivation for alcohol following three weeks after the SD procedure resumed.

The eCS plays a fundamental role in drug-related behaviors and the stress response (Lutz et al., 2015; Sagheddu et al., 2015). The upregulation of eCB and eCB-like lipids after alcohol or stress exposure has been suggested to represent a natural defense mechanism (Pertwee, 2014). According to these observations, we hypothesized that exogenous administration of OEA closely to the stress episode could strengthen the neuroprotective effects of eCS.

Previous studies have provided evidence for an attenuating effect of OEA on alcohol drinking (Bilbao et al., 2016; Shahen-Zoabi et al., 2023; González-Portilla, *in preparation*). Nevertheless, to our knowledge, this is the first study to explore the effects of OEA on stressed-related alcohol SA. Our data demonstrates that OEA (10mg/kg) was effective in reducing stress-induced alcohol consumption when administered chronically (10 daily doses) after the SD protocol (EXP/SD-10OEA groups). These results are consistent with a recent study in which daily OEA treatment (10mg/kg) for 14 days prevented the behavioral alterations induced by chronic SD (Rani et al., 2021).

In addition, in the EXP/SD-OEA group we administered OEA 15 minutes before the SD to test whether OEA could function as a protection for stress-induced drinking. However, in this OEA-treated group, alcohol drinking nor active nose poke responses did not differ from that of non-treated SD mice. It is possible that a higher number of doses are required to observe a long-lasting effect that more robustly diminish alcohol consumption.

In contrast to the results from stress-induced cocaine CPP (González-Portilla et al., 2022), we observed that chronic and not contingent acute treatment before SD buffered the stress-induced vulnerability to the rewarding properties of drugs of abuse. These diverging results may result from the pharmacological specificity of the drug employed (cocaine vs. alcohol) and the dose chosen. In the previous study, CPP was conducted with a subthreshold cocaine dose while the SA was performed with an effective alcohol solution 20% (vol/vol), which may reflect different aspects of drug reward. In addition, discrepancies between results may stem from the different tests employed. The CPP assesses the strength of drug-associated memories which throws a measure of drug reward. In the SA, instead, as an operant-based procedure, drug delivery is a measure of both drug reward and motivation. As two fundamentally different paradigms, performance largely also rely on different brain substrates (Spanagel, 2017).

A hallmark of AUD is the recurring relapse episodes which may has been associated to an increased cue reactivity (Wilcox et al., 2011; Zeng et al., 2021).Other studies have reported that OEA administration previous to the cue-induced reinstatement test can prevent relapse (Bilbao et al., 2016; González-Portilla et al., in preparation). Alternatively, we tested the long-term effects of OEA on counteracting the stress effects on the susceptibility to reinstate alcohol seeking. Our data shows that both EXP and SD mice reinstated alcohol seeking when the context cues become available. However, we did not find any influence of OEA on reinstatement.

The mechanisms of OEA action on alcohol SA remain to be elucidated. The main molecular target of OEA is peroxisome proliferator receptor alpha PPAR-α, a transcription factor which regulates lipid metabolism and inflammation pathways (Wahli & Michalik, 2012). The molecular action of PPAR-α is responsible for many of the physiological effects of OEA, including its effects on alcohol drinking (Kaczocha et al., 2012). In this sense, administration of OEA in PPAR-α knock-out mice failed to reduce voluntary alcohol drinking and prevent cue-induced reinstatement of alcohol-relapse. Similarly, pharmacologically antagonizing PPAR-α with GW6471, the beneficial effects of OEA were abolished (Bilbao et al., 2016).

Our results suggest that boosting OEA levels after SD exposure may prevent the adverse effects of stress, possibly by its antidepressant activity. Previous work in our laboratory suggests that OEA inhibits inflammatory pathways involved in social stress (González-Portilla et al., 2022). According to these evidence, increased OEA levels may buffer the immune adverse effects of SD.

In conclusion, our findings reveal that systemic OEA administration diminished SD-induced increase in alcohol consumption in the SA paradigm. The reduction in alcohol drinking was only observed with a chronic schedule after SD and not with an acute contingent treatment. Nevertheless, OEA treatment did not affect behavior on cue-induced reinstatement. Taken together, these results suggest that boosting OEA levels may be an effective strategy to mitigate some of the adverse effects of social stress.

